# Cell death in cells overlying lateral root primordia contributes to organ growth in *Arabidopsis*

**DOI:** 10.1101/268433

**Authors:** Sacha Escamez, Benjamin Bollhöner, Hardy Hall, Domenique André, Béatrice Berthet, Ute Voß, Amnon Lers, Alexis Maizel, Malcolm Bennett, Hannele Tuominen

## Abstract

Unlike animal development, plant organ growth is widely accepted to be determined by cell division without any contribution of cell elimination. We investigated this paradigm during *Arabidopsis* lateral root formation when growth of the new primordia (LRP) from pericycle-derived stem cells deep inside the root is reportedly facilitated by remodeling of the walls of overlying cells without apparent cell death. However, we observed the induction of marker genes for cell types undergoing developmental cell death in several cells overlying the growing LRP. Transmission electron microscopy, time-lapse confocal and light sheet microscopy techniques were used to establish that cell death occurred at least in a subset of endodermal LRP-overlying cells during organ emergence. Significantly, organ emergence was retarded in mutants lacking a positive cell death regulator, and restored by inducing cell death in cells overlying LRP. Hence, we conclude that in the case of LRP, cell elimination contributes to organ growth.

---

In contrast with the regulation of animal organ growth^1^, cell elimination is generally considered not to play a role in regulating plant organ growth^2-4^. Nevertheless, the growth of the embryo in the seed of the model plant *Arabidopsis thaliana* (hereafter *Arabidopsis*) is associated with the elimination of the neighboring endosperm tissue^5^. In addition, the growth of the root organ in *Arabidopsis* depends on cell elimination in the lateral root cap which surrounds the root tip^6^. Several types of cells die and degrade their own protoplast by autolysis as part of plant development, a phenomenon referred to as programmed cell death (PCD)^7-10^ which implies a mechanism of cell suicide^11-13^. However, in cases such as the *Arabidopsis* endosperm the PCD framework does not apply because the death of these cells is caused by the growing embryo^5^. Similarly, loss of viability in cells next to growing plant organs may not only be due to endogenous (i.e. PCD) but also exogenous mechanisms. It is therefore better to describe the loss of viability and the subsequent autolysis in connection to the plant organ growth simply as “cell death” to avoid implicit mechanistic assumptions. It remains to be investigated whether such cell death events contribute, in addition to cell proliferation, to growth of neighboring organs more widely in plants.

Features of cell death and autolysis have been observed in several species in cells overlying the sites of lateral root (LR) formation within existing roots^14-17^. The lateral roots are initiated from a subset of pericycle cells which form the lateral root primordium (LRP) deep in the parent organ^18, 19^. The developing LRP must therefore traverse the overlying endodermal, cortical, and epidermal cell layers for lateral root emergence (LRE) to occur. LRE has been shown to rely on cell divisions and turgor-driven expansion in the LRP^18, 20^, but also on changes in the cell walls and shapes of the LRP-overlying cells^20-26^. Cell death is not believed to happen during LRE in *Arabidopsis*^27^ and the cell death reported in the LRP-overlying cells of other species^14-17^ has not been studied in relation to LRP growth, leaving the question of whether cell death contributes to LRE open.

In *Arabidopsis*, the most dramatic changes reported during LRE occur in the LRP-overlying endodermal cells due to their position in immediate contact with the LRP and the presence of their lignified casparian strip cell wall region. In front of the growing LRP, endodermal cells modify their shape to such an extent that they occasionally split, with both halves having the ability to maintain their integrity at least for some time^25^. The cortical and epidermal cells are less affected, as their cell walls are loosened by hydrolytic enzymes so that they can separate to allow the emerging LRP to pass through^21, 27^. There is currently no report of cell death during LRE in *Arabidopsis*, suggesting that the remodeling and separation of the LRP-overlying cells is sufficient to ensure LRE without any contribution from cell death. Yet, cysteine proteases associated with cell death and autolysis are expressed in LRP-overlying cells^28^, supporting the occurrence of cell death during LRE in *Arabidopsis*.

The present study revisits whether cell death occurs in the LRP-overlying cells during LRE in *Arabidopsis*, and if so, whether it contributes to LRP growth leading to LRE. We detected expression of several canonical marker genes for cell types undergoing developmental cell death^29^ in a subset of LRP-overlying cells. Electron microscopy revealed autolytic features indicative of cell death in some of the endodermal cells overlying early-stage LRPs. Live cell imaging by confocal and light sheet microscopy confirmed that cell death occurred in a subset of the LRP-overlying endodermal cells, concomitant with the growth of the LRP through the endodermis. Plants unable to express ORESARA1/ANAC092 (ORE1), a transcription factor known to activate the expression of several cell death related genes^30^, displayed a delay in LRE. When cell death was restored in the overlying cells of these plants by expressing the mammalian cell death promoting factor mBAX^31-33^ LRP growth reverted to normal, indicating that cell death contributes to regulating organ growth during LRE.

## Results

### Cell death indicator genes are induced in cells overlying lateral root primordia

In a time-course transcriptomics dataset covering various stages of LRP growth^34^ we detected upregulation of *BIFUNCTIONAL NUCLEASE 1* (*BFN1*) which functions in cell autolysis associated with developmental cell death^6^ (Table S1). Several other genes expressed in cell types undergoing developmental cell death and autolysis^29^ were identified among the genes most correlated with *BFN1* in the LRE transcriptome. Among them, *METACASPASE 9* (*MC9*), *RIBONUCLEASE 3* (*RNS3*), *EXITUS 1* (*EXI1*) and *DUF679 DOMAIN MEMBRANE PROTEIN 4* (*DMP4*) together with *BFN1* represent five out of the nine core marker genes specifically expressed in cell types undergoing developmental cell death in *Arabidopsis*^29^ and are thereafter referred to as “cell death indicator genes” (Table S1).

The five cell death indicator genes displayed three peaks of expression in the LRE transcriptome^34^, coinciding temporally with the passage of the growing LRP through each of the three overlying (endodermal, cortical and epidermal) cell layers (Fig. 1a). Using promoter::*GUS* reporter lines, we also detected activation of the promoters of the cell death indicator genes in LRP-overlying cells at different stages of LRP growth (Fig. 1b), defined according to Malamy and Benfey^18^. A more detailed time-lapse confocal microscopy analysis of a pro*BFN1::*nuc*GFP* reporter line^35^ carrying the pro*UB10::WAVE131*:*YFP* plasma membrane marker^36^ revealed the activity of the *BFN1* promoter in just a few endodermal cells overlying an early stage LRP, as well as in overlying epidermal cells (Fig. S1).

**Figure 1:**
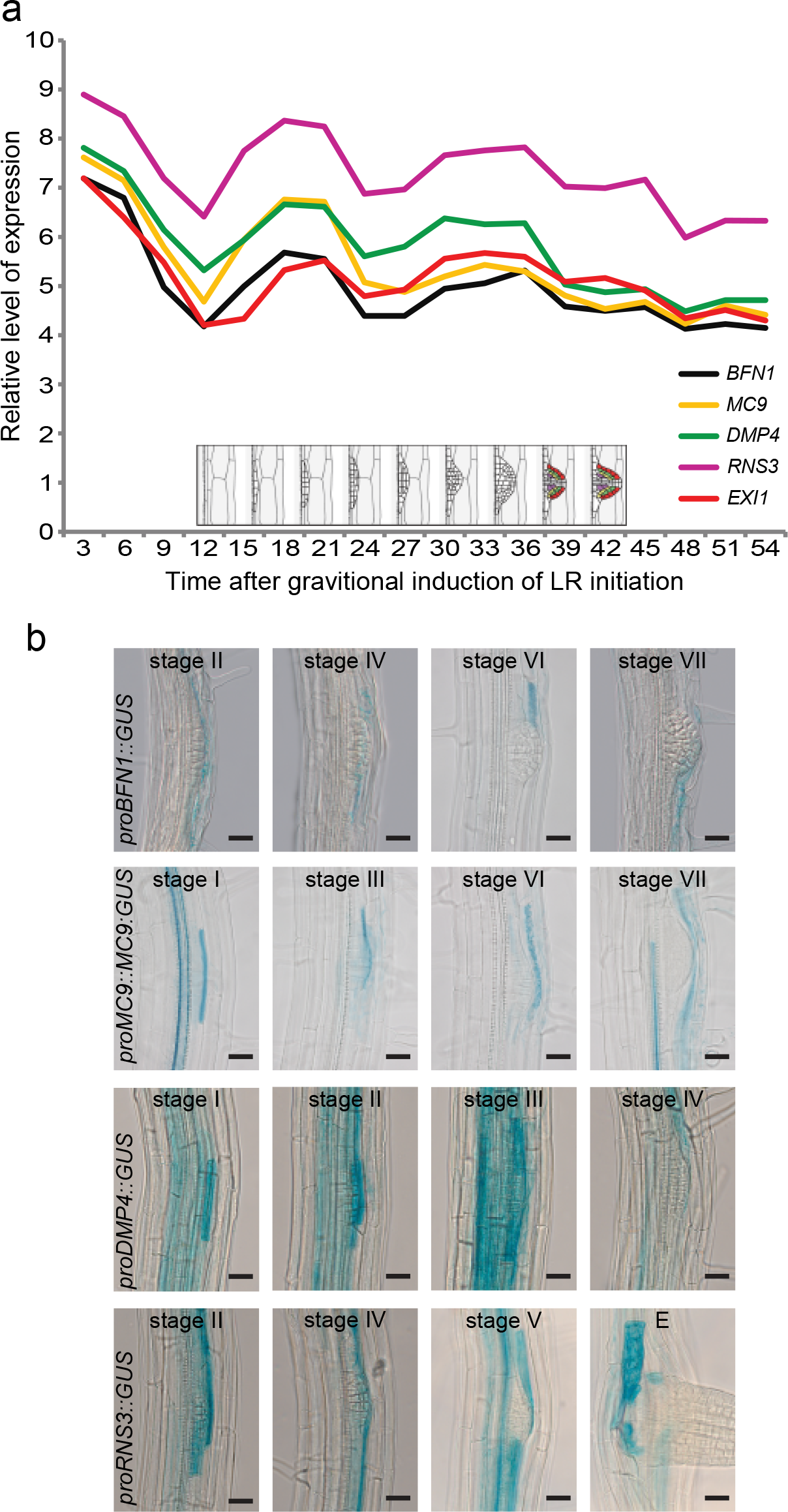
Transcriptional activation of cell death indicator genes in LRP-overlying cells. (a) Expression profile of *BFN1* and four highly correlated cell death indicator genes in the lateral root (LR) emergence time course transcriptomics dataset from ref 34. (b) Promoter activity profile of the cell death indicator genes *BFN1*, *MC9*, *DMP4* and *RNS3* in the tissues overlying naturally initiated lateral root primordia (LRP) at the indicated stages. Bars = 25 μm. Note that in addition to signal in the overlying cells there is also often signal in the protoxylem vessel.

Next, we assessed the relationship between transcriptional induction of the cell death indicator genes and the LRP growth by calculating the proportion of LRP with pro*BNF1::*nuc*GFP* signal in the LRP adjacent cells that were either overlying the LRP (during early stages of LRP growth) or neighbouring the LRP (in late stages of growth) (Fig. 2a,b). A total of 204 (53.5 %) out of the 381 observed loci with LRP displayed pro*BNF1::*nuc*GFP* signal in at least one adjacent endodermal cell, while 56 (14.7 %) and 37 (9.7 %) showed signal in at least one cortical or epidermal cell, respectively. The proportion of LRP with pro*BNF1::*nuc*GFP* signal in at least one adjacent cell was even higher when calculated in relation to the specific stage of LRP development (Fig. 2a). In the endodermis, the frequency of pro*BNF1::*nuc*GFP* signal adjacent to an LRP reached its maximum (over 90%) at stage IV, while the cortex displayed signal most frequently (nearly 50 %) at stage VII and the epidermis (40%) at stage VIII (Fig. 2a). Interestingly, we detected lower frequency of pro*BNF1::*nuc*GFP* signal in cells adjacent to LRP whose growth was delayed (Fig. 2b). Taken together, these results indicate that cell death indicator genes are almost always induced in at least one LRP adjacent endodermal cell and also, however less frequently, in cortical and epidermal cells, in connection with LRP growth.

**Figure 2:**
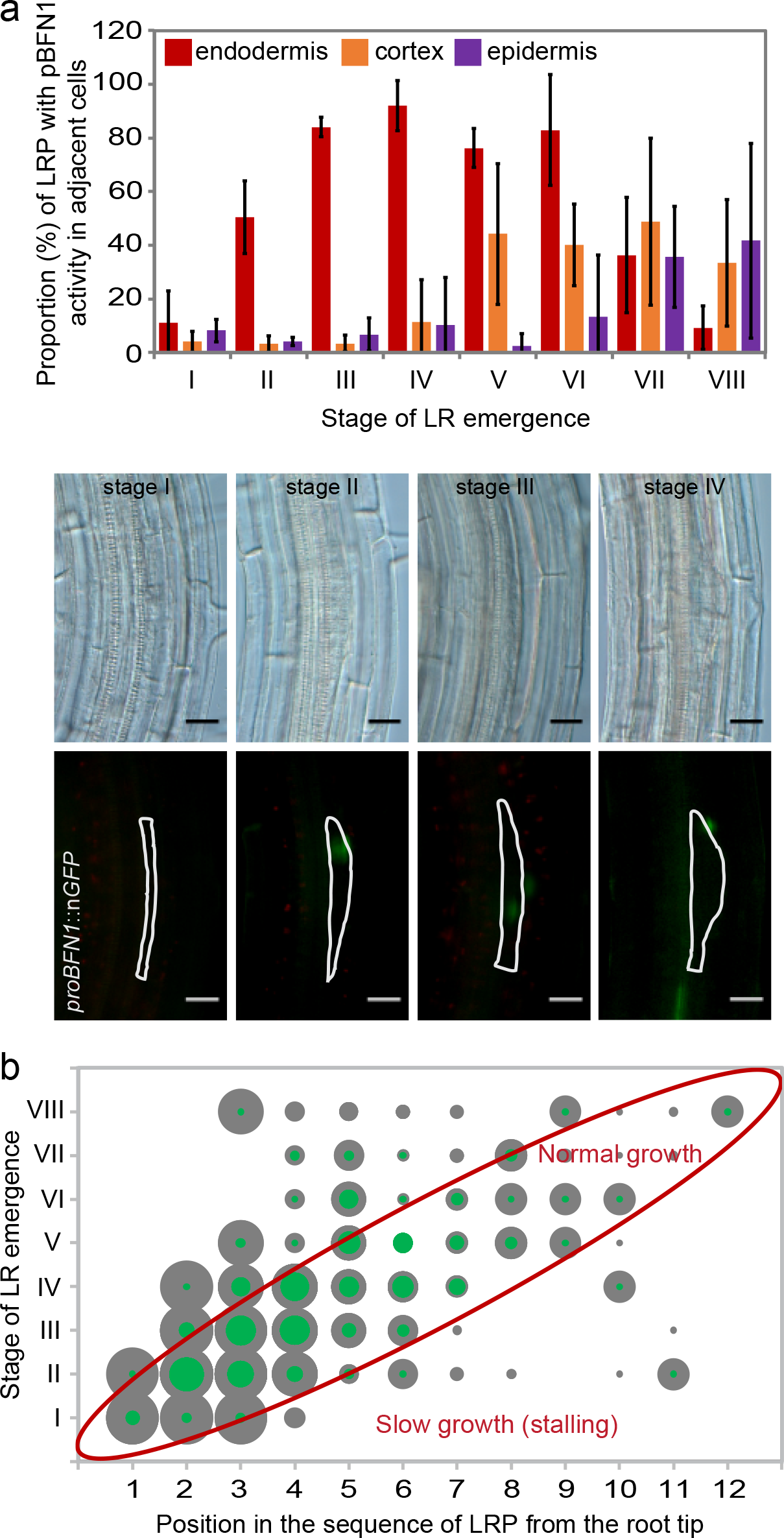
Expression patterns of cell death indicator genes in LRP-overlying cells in relation to LRP progression. (a) Proportion of LR primordium (LRP) displaying nuclear GFP signal in at least one LRP adjacent endodermal cell (red), cortical cell (orange), or epidermal cell (purple) for each stage of LRP development in the pro*BFN1*::nuc*GFP* seedling roots. Values represent averages of three replicate experiments, each including 20 seedlings and over 100 primordia. Error bars indicate SD. The micrographs in the lower panels illustrate how the data was acquired by differential interference contrast microscopy (upper row) for analysis of the LRP stage and by fluorescence microscopy (lower row) for GFP signal count in the marker line pro*BFN1*::nuc*GFP*. The LRP shape is delineated by a white line in the fluorescence micrographs. Bars = 25 μm. (b) Visualization of LRP stage distribution in terms of LRP sequence from the root tip of 60 5-6 days old Arabidopsis seedlings. The size of each dot is proportional to the number of observed LRP at each stage and position in total (grey) or with at least one pro*BFN1::*nuc*GFP*-positive overlying cell (green). E = emerged.

### Cell death occurs in a subset of LRP-overlying cells

To determine the fate of the cells overlying the growing LRP, we employed several cell death detection methods. Terminal deoxynucleotidyl transferase dUTP nick end labeling (TUNEL), marking the DNA strand breaks which follow cell death, could be detected in the nuclei of cells close to the growing LRP (Fig. 3a). Hence, TUNEL revealed that a subset of LRP-overlying cells was undergoing autolytic processes associated with cell death.

**Figure 3:**
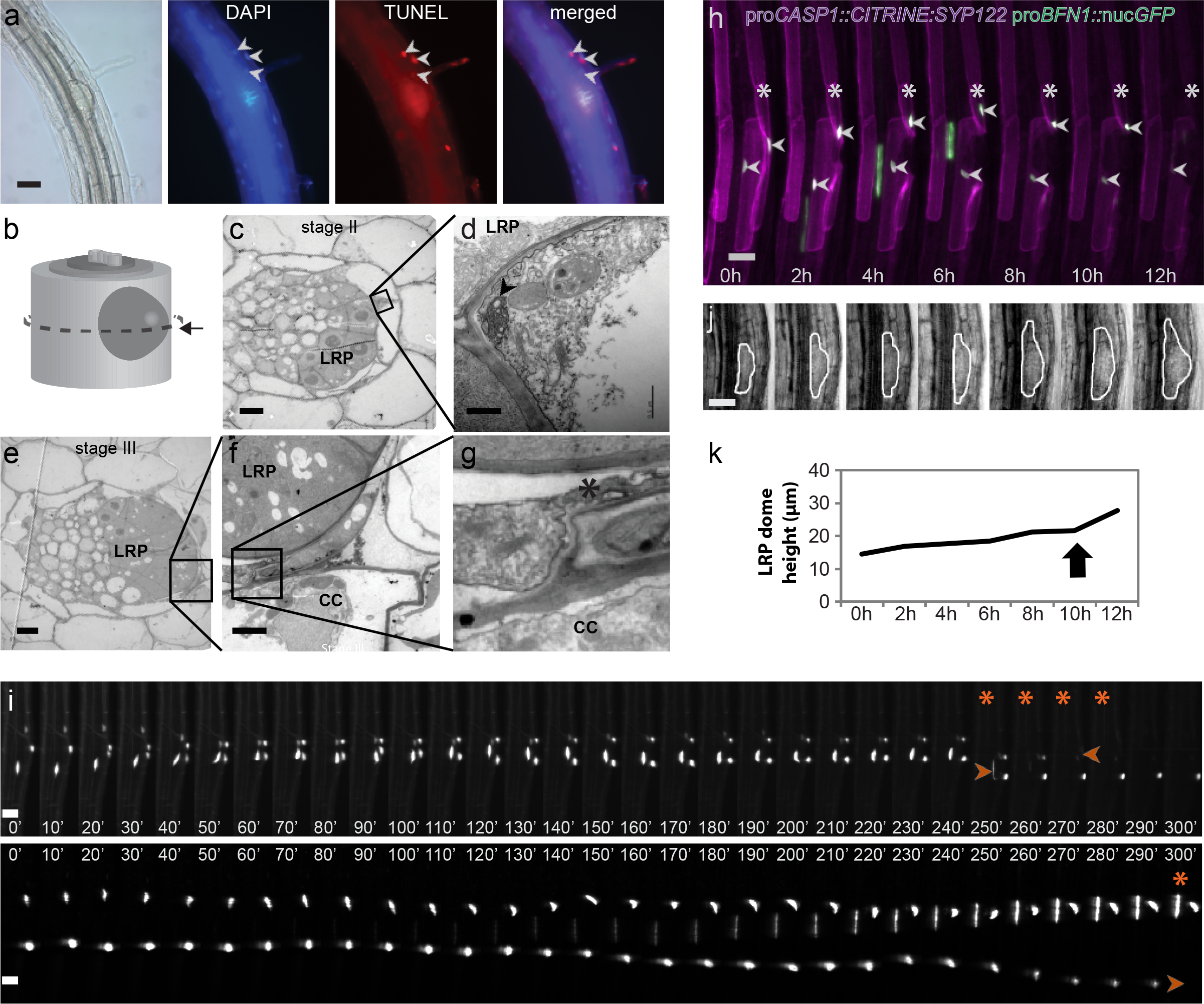
Detection of cell death in LRP-overlying cells. (a) Fluorescence micrography detection of cell death-associated nuclear degradation with TUNEL (red) and ubiquitous nuclear staining with DAPI (blue). Arrowheads indicate TUNEL-positive nuclei in LRP-overlying cells. Bar = 50μm. (b-g) Transmission electron micrography of cross sections through a lateral root primordium (LRP) and the surrounding tissues. The position of the cross section within the root is shown in (b). Stage II (c,d) and stage III (e-g) LRPs are shown. (d) is a magnification from (c). (g) is a magnification of (f) which itself is a magnification of (e). The arrowhead indicates apparent loss of plasma membrane integrity while the asterisk indicates leakage of intracellular material. LRP, lateral root primordium; CC, cortical cell. Bar = 5 μm (c,e), 2 μm (f) and 0.5 μm (d). (h) Time-lapse confocal microscopy 3D projection of the endodermal plasma membrane marker pro*CASP1*::*CIT:SYP122* (purple) and the nuclear cell death reporter pro*BFN1*::nuc*GFP* (green) around a developing LRP. The asterisks mark the endodermal cell which ultimately dies (between 10h and 12h) while the arrowheads point at the endodermal nuclei displaying GFP signal. The elongated area with GFP signal seen in the stele from 2h to 6h is the nucleus of a xylem tracheary element undergoing developmental cell death and autolysis. Bar = 25μm. (i) Kymographs of 3D projection from light sheet microscopy time-lapse imaging of the endodermal plasma membrane marker pro*CASP1*::*CIT:SYP122* and the nuclear-localized cell death indicator gene transcriptional reporter pro*BFN1*::nuc*GFP* around a developing LRP. The presence of a LRP can be deduced from the bent shape of the overlying endodermal cells. Asterisks indicate time points when an endodermal nucleus is seen to disintegrate or to disappear. Arrowheads point at disintegrating nuclei. Bars = 20 μm. (j) Transmission light images corresponding to images in (h). The LRP are highlighted by white lines. Bars = 25 μm. (k) Dome height (in μm) of the LRP from (h,j) over the 12 h monitoring period of LRE. The black arrow indicates acceleration of LRP growth along its length axis concomitant with the death of an overlying cell.

The LRP-overlying endodermal cells displayed transcriptional activation of the cell death indicator genes more frequently than other cell types (Fig. 2a). Because endodermal cells undergo more dramatic shape changes than other overlying cell types during LRP development^25^, we studied the sub-cellular morphology of these cells in more detail by employing transmission electron microscopy (TEM) on root cross sections containing LRP at early stages (Fig. 3b). Plasmolysis and autolytic features indicative of cell death, such as loss of plasma membrane integrity and leakage of intracellular material outside of the protoplast, could be observed specifically in endodermal cells overlying early stage LRP (Fig. 3c-g). These TEM observations are therefore consistent with the occurrence of cell death and autolysis in a subset of LRP-overlying endodermal cells.

Endodermal cells were also observed by time-lapse confocal and light sheet microscopy using the endodermis-specific plasma membrane marker pro*CASP1::CITRINE:SYP122*^25^. We simultaneously relied on the pro*BFN1::*nuc*GFP* marker which not only revealed transcriptional activation of cell death related genes but also cell death itself, based on the abrupt disappearance of nuclear GFP signal known to shortly follow cell death^6, 35^. Time-lapse confocal microscopy imaging of LRP provided evidence for the occurrence of cell death in endodermal cells overlying LRP in two out of five cases, as revealed by complete loss of nuclear GFP signal in these endodermal cells between two consecutive time points (Fig. 3h; Fig. S2). Light sheet microscopy (which provided a 12-fold greater time resolution) revealed complete disappearance of nuclear GFP signal between two consecutive time points in the LRP-overlying endodermal cells of two seedlings (Fig. 3i; Movies S1,2). One of these endodermal nuclei disintegrated just before losing its signal (Fig. 3i; Movies S1). Both the apparent nuclear disintegration and the rapidity of the nuclear signal disappearance can only be explained by cell death. The shapes of the plasma membranes of the LRP-overlying cells at the times of death indicated that the LRP had not yet entirely traversed the endodermis (Fig. 3h,i; Fig. S2; Movie S3), meaning that the observed cell death events occurred either before or during the passage of the LRP through the endodermal layer. A number of surviving endodermal cells in close proximity to LRP displayed a slow and gradual decrease of the nuclear GFP signal (Fig. 3h,i; Fig. S2), suggesting deactivation of the cell death and autolysis transcriptional machinery in these cells. A few other endodermal cells kept a high level of nuclear GFP signal over the observation time span (Fig. 3i; Fig. S2) and it cannot be concluded whether or not these cells would have died at a later point during LRP emergence.

The height of the dome shape of five LRP observed with time-lapse confocal microscopy was measured at the same time points at which the endodermal nuclear GFP fluorescence was recorded, revealing accelerations of LRP growth concomitant with the death of an overlying endodermal cell (Fig. 3j,k; Fig. S2). Albeit weakly, LRP growth was on average better correlated with the dynamics of nuclear GFP signal intensity of the overlying endodermal cells that died (squared Pearson correlation coefficient r^2^=0.32) than with the nuclear signal dynamics of the non-dying cells (r^2^=0.03), suggesting causality between LRP progression and endodermal cell death.

### Cell death in LRP-overlying cells facilitates LRP growth

We reasoned that if cell death played a role in facilitating LRP growth, then plants impaired in parts of the cell death machinery may show delayed LRE. To compare the speed of LRE between genotypes, LR initiation was induced synchronously by gravitational stimulus (90° rotation of the seedlings)^18, 20^. When monitored at 18h and 42h post-gravitational induction (pgi), the single mutants for the cell death indicator genes did not show any consistent or significant changes in LRP growth (Fig. S3), possibly because of functional redundancy.

A large number of cell death related genes, including the five cell death indicators identified in this study, are transcriptionally regulated by the NAC transcription factor ORESARA1/ANAC092 (ORE1)^30^. ORE1 is expressed in connection with several types of developmental cell death and autolysis^37^ and is also connected to LR development^38^. We therefore hypothesized that ORE1 might contribute to the transcriptional control of cell death in the LRP-overlying cells, and that analyzing LRP growth in the *ore1* mutants could overcome potential genetic redundancies between cell death related genes during LRE. Consistent with a role for ORE1 during LRE, shorter LRs were observed in two *ore1* mutant alleles compared to wild-type seedlings (Fig. S4a-c). Gravitational induction experiments also revealed delayed LRP growth in these *ore1* mutants (Fig. 4a).

**Figure 4:**
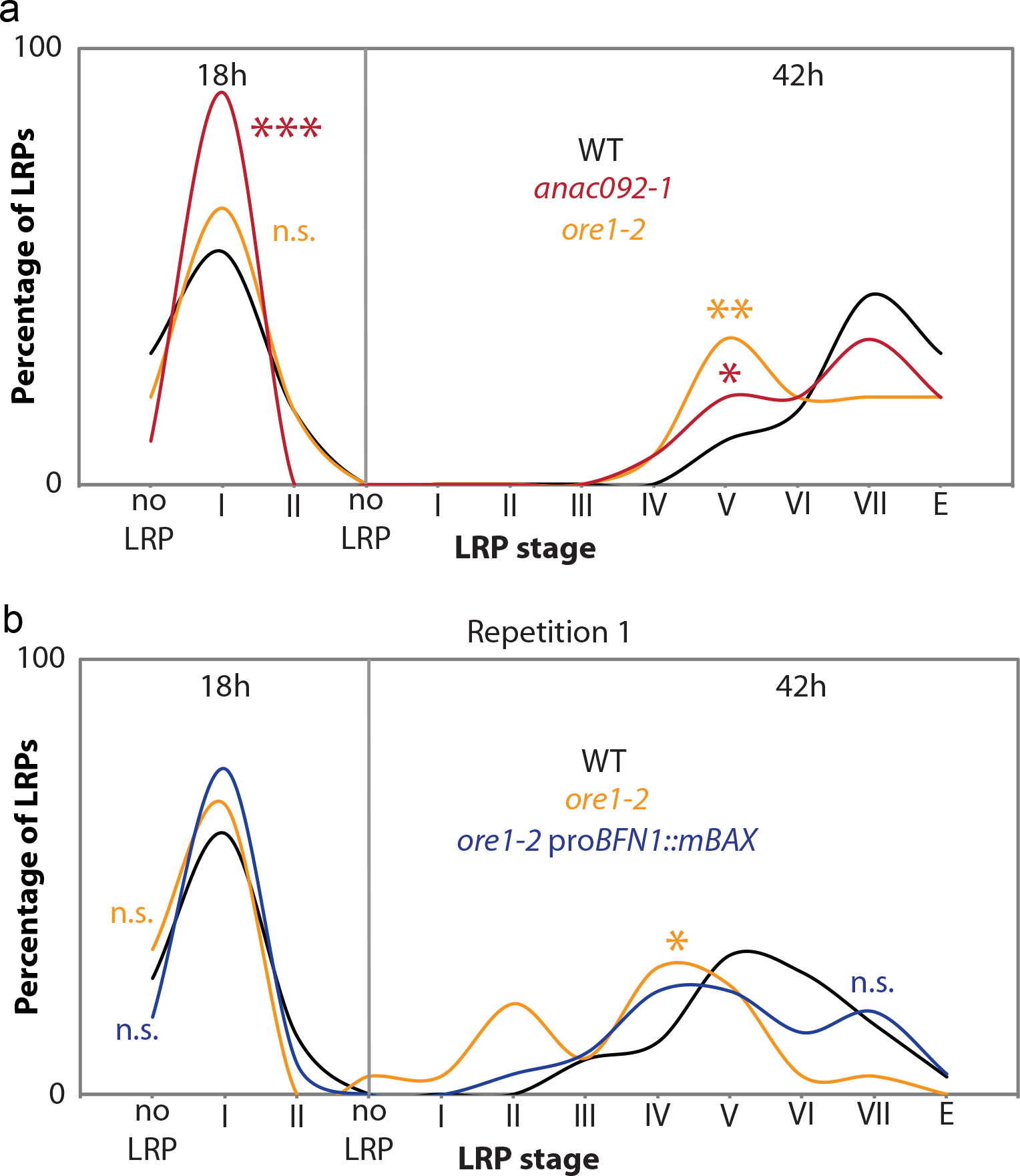
LRE is delayed by loss of the positive cell death regulator ORE1 and rescued by inducing cell death in overlying cells. (a) Distribution of observed lateral root primordium (LRP) stages 18h and 42h after gravitational induction of lateral root (LR) initiation in wild type (WT) and two mutant alleles for ORE1. (b) Distribution of observed LRP stages 18h and 42h after gravitational induction of LR initiation in WT, *ore1-2* mutant and *ore1-2* expressing the cell death inducing mBAX protein under the transcriptional control of the *BFN1* promoter. Each line was compared with the corresponding wild type by Pearson’s chi-square test, to reveal potential differences in the distribution of LRP stages dependent on genotype (n.s.: not significant, *: p<0.05, **: p<0.01, ***:<0.001). In (b), 20<n<31, while in (a), n=30 observed seedlings.

To test whether *ore1* mutants displayed slower LRP growth as a result of altered cell death in the overlying cells, we set out to rescue the LRE delay of *ore1* by inducing cell death in cells overlying LRP. To do this, we expressed the pro-apoptotic mammalian protein mBAX^31^, known to potently induce cell death in *Arabidopsis*^32, 33^, under the transcriptional control of the *BFN1* promoter (pro*BFN1::mBAX*) in the *ore1-2* background. This line expressed mBAX (Fig. S4d) and showed signs of cell death in front of LRP (Fig. S5), but did not display obvious alterations in plant development most probably because the *BFN1* promoter is active only at a late stage of development in cells that are anyway destined to die^29^.

Interestingly, the LRP growth was restored to at least wild-type level in the *ore1-2* mutants expressing *mBAX* under the transcriptional control of the *BFN1* promoter (Fig. 4b; Fig. S4b,c,f). The fact that the cell death inducing pro*BFN1::mBAX* construct had a rescuing effect on LRE in *ore1-2* (Fig. 4b; Fig. S4f) but no effect in the wild-type plants (Fig. S4e) indicates that ORE1 regulates cell death which normally occurs in cells overlying LRPs. Collectively, the correlation between LRP growth dynamics and overlying cell death (Fig. S2), and the rescue of LRP growth in *ore1-2* by introducing expression of mBAX show that cell elimination contributes to regulating LRP growth.

## Discussion

Our study demonstrates that, contrary to what was previously thought, cell death occurs in cells overlying LRP in *Arabidopsis* and also contributes to regulating LRP growth. The occurrence and relevance of cell elimination during LRE of *Arabidopsis* is likely to have been previously overlooked for several reasons. First, it is clear that living LRP-overlying cells undergo a number of major changes which contribute to LRE^20-26^, rendering the occurrence of cell death seemingly unnecessary. Second, cell death has not been observed during LRE in *Arabidopsis* probably because following cell death of a few cells among a large number of viable cells is difficult unless high-resolution methods and suitable markers are used. Monitoring of plasma membrane or tonoplast integrity by confocal microscopy of membrane markers or propidium iodide staining^25^ may not have provided the necessary resolution and contrast to unequivocally detect cell death of these few cells. Furthermore, the mechanism of cell death in LRP-overlying cells is not known and may differ from other instances of developmental cell elimination. This is supported by the fact that only five of the nine canonical *Arabidopsis* developmental cell death indicator genes^29^ were highly co-expressed during LRE (Table S1; Fig. 1a). In addition, expression of a tonoplast integrity marker^6^ under the control of the *BFN1* promoter revealed a longer time gap between transcriptional activation of this promoter and loss of tonoplast integrity in LRP-overlying cells than in xylem tracheary elements (Movie S4) which undergo a well-studied case of developmental cell death and autolysis^39^. Moreover, a lag of several hours could be observed with the tonoplast integrity marker^6^ between the loss of tonoplast integrity and the disappearance of nuclear signal in LRP-overlying endodermal cells (Movie S5), unlike in other dying cell types^6, 40^. Finally, if one carefully looks at published images of emerging lateral roots, it is possible to distinguish cells whose collapsed morphology is suggestive of their death (see for instance Figure 1c in^41^). Hence, our observations are not necessarily in conflict with earlier observations in *Arabidopsis*, and are further supported by observations of the demise of cells overlying LRP in other species^14-17^.

The spatial and temporal pattern of cell death progression during LRE suggests the existence of a signal responsible for activating the genetic modules of cell elimination. Both LRP growth and remodeling of LRP-overlying cells rely to a significant extent on the hormone auxin^20, 23, 24^. Auxin also accumulates in several cell types that ultimately undergo developmental cell death^42, 43^. However, the cell death and autolysis related genes studied here are not known to be induced by auxin except for *ORE1*^38^. It is therefore conceivable that auxin triggers a signaling cascade towards cell death and autolysis in front of the LRP through induction of upstream regulators such as ORE1. On the other hand, ORE1 does not appear to be a primary auxin response gene as its induction by this hormone is slower than by abscisic acid (ABA) or the ethylene precursor ACC^38^. Hence, we propose that the induction of cell death and autolysis in the LRP-overlying cells involves primarily other signals than auxin. Credible candidates include ABA and ethylene.

Mechanical forces are also expected to influence the fate of the LRP-overlying cells. Strong mechanical pressure is likely to be exerted by the LRP growing against the overlying cells which themselves alter their mechanical properties and rigidity in coordination with LRP growth^20, 22^. This could eventually result in the death of some weakened overlying cells under intense pressure. Such a scenario is analogous to the elimination of *Arabidopsis* endosperm cells which, by loosening their cell walls, allow the embryo to grow into the space that they occupy and then mechanically kill them^5^. Similarly, during adventitious root formation, mechanical pressure has been reported to induce cell death in the epidermis of rice in an ethylene-dependent manner^44^. It is therefore possible that mechanical pressure represents at least one of the mechanisms involved in cell death signaling, or even in the execution of cell death itself, during LRP growth. Furthermore, such LRP-mediated mechanical pressure would likely affect the overlying cells differently depending on their own mechanical properties and on their degree of overlap with the growing LRP, thereby providing a possible explanation for the fact that only a subset of the LRP-overlying cells dies.

We furthermore demonstrated that cell death contributes, along with several other processes^20-26^, to the growth of the LRP. Even though the underlying mechanism remains to be tested, it is likely that cell death facilitates growth of the LRP by reducing the mechanical resistance of the overlying cells towards the primordia. The plausibility of such mode of action based on mechanical feedback is supported by the fact that laser-ablation of an endodermal cell is potent enough to trigger LR initiation in the adjacent pericycle^45^, likely because the loss of mechanical feedback from the ablated cell allows the necessary swelling of the pericycle to initiate LRP formation^25, 46^. Alternatively, or in addition, cell death might contribute to cell wall remodeling and cell separation and hence to LRP growth by allowing a massive release of cell wall modifying enzymes and/or of auxin. For example, cell death of a few columella stem cell daughter cells in *Arabidopsis* roots exposed to low temperature was recently shown to affect auxin distribution^47^. Furthermore, the developmental cell death of lateral root cap (LRC) cells was shown to result in release of auxin that is necessary for LR initiation^43^. Interestingly, LRP growth was delayed to a comparable extent in mutants for ORE1 and for the LATERAL ORGAN BOUNDARIES-DOMAIN/ASYMMETRIC LEAVES 2-LIKE29 (LBD29) transcription factor, which normally controls auxin accumulation in LRP-overlying cells^23^ (Fig. 4; Fig. S4a). It is therefore tempting to hypothesize that the death of LRP-overlying cells may serve an analogous purpose to LRC cell death in quickly releasing high amounts of auxin for cell wall remodeling of other overlying cells and for LRP growth.

Our finding on the impact of cell death on lateral root growth demonstrates that plant organ growth can be regulated by cell elimination. It was also recently shown that embryo growth requires cell death of the bordering endosperm^5^ and that root organ growth is regulated by cell death in a cell type which borders its tip^6^. These findings therefore call for a major shift in opinion to accept that cell proliferation is not the sole factor determining organ growth in plants and that the regulation of organ growth in plants is not as dissimilar from that in animals as previously proposed.

This greater similarity than previously accepted in the regulation of organ growth between animals and plants raises some evolutionary questions. Has cell elimination arisen as a mechanism to regulate organ growth from a shared evolutionary heritage between animals and plants, for example from the regulation of early unicellular populations by cell death^8^? Alternatively, could the regulation of organ growth by cell death in animals and plants have stricken roots at different locations in the tree of life, making it a form of convergent evolution which may reveal in the future some form of deep evolutionary constraint linked with cell elimination?

## Acknowledgements

We thank Joop Vermeer for the pro*CASP1::CITRINE:SYP122* line, Moritz Nowack for the tonoplast integrity marker, Laszlo Bako for the proLBD16::GFP line, Michael Wilson and Kim Kenobi for bioinformatic assistance, Kamal Swarup and Julien Lavenus with confocal imaging, and Lenore Johansson from the Umeå Core Facility for Electron Microscopy (UCEM) for technical assistance. This work was supported by the Swedish Research Council VR (621-2013-4949), the Swedish Research Council Formas (232-2009-1698), and the Swedish Governmental Agency for Innovation Systems VINNOVA (015-02290) as well as by the Boehringer Ingelheim Fundation and Deutsch Forschung Gemeinschaft (FOR2581) and Landesgraduiertenförderung to BBe and AM.

## Author contributions

H.T. and M.B. conceived the study. S.E., B.Bo., H.H., D.A., and B.Be. performed the experiments, A.M., M.B., S.E., B.Bo. and H.T. supervised the experiments, and S.E. and H.T. wrote the manuscript.

## Material and Methods

### Plant Material and Growth conditions

Most of the *Arabidopsis* plants used in this study are in the Columbia-0 (Col-0) genetic background, and were therefore compared with a Col-0 wild type: *anac092-1* (*ore1* allele SALK_090154^30^), *ore1-2* (5bp deletion^48^), *bfn1-1* (GK-197G12^6^), *bfn1-2* (SALK_017287), *mc9-1* (GABI_540_H06^35^), *mc9-2* (SALK_075814^35^), SALK_063946C (for *DMP4*), SAIL_195_E01 (for *RNS3*), SAIL_760_G07 and SALK_137383C (for *EXI1*). The FLAG_164A04 allele for *RNS3* was in the Wassilewskija (Ws) genetic background, and compared to the corresponding wild type.

To compare the speed of LRE between genotypes, seedlings were grown on ½ MS medium for three days in constant light (150 μE·m^−2^·s^−1^) on vertical plates, which were then rotated 90° to induce LRP initiation^20^. Determination of LRP stage was performed 18h and 42h after synchronized LR induction by gravitational stimulus on cleared samples observed as described previously^18, 20^.

Seedlings whose naturally initiated LRPs (no induction) were observed with any of the presented microscopy techniques were grown on ½ MS medium for 5 - 7 days in 16 h day (150 μE·m^−2^·s^−1^) / 8 h night cycles.

### *BFN1* co-expression analysis

To identify genes correlating in expression with *BFN1* during LR development, smooth splines were fitted through the transcriptomic profiles for all genes, to smooth out the noise. The Pearson correlation between each gene’s profile and that of the target gene *BFN1* was calculated, including all data points, i.e. not just mean values for each time point. Genes were ranked in decreasing order of the Pearson correlation with *BFN1*.

### Cloning and plant transformation

For promoter-reporter constructs, 1654 bp (AT4G18425; *DMP4*) or 2000 bp (AT1G26820; *RNS3*) upstream of the translational start codon were amplified and recombined by Gateway-mediated cloning *via* pENTR207 (Invitrogen) into the destination vectors pBGGUS^49^. Col-0 plants were transformed with *Agrobacterium tumefaciens* by floral dip as previously described^50^. At least five independent lines were analysed for each construct to select representative lines. pro*BFN1::GUS* was obtained by recombining a 670 bp promoter region of *BFN1* into pMDC163. pro*BFN1::*nuc*GFP*, pro*MC9::*nuc*GFP* and pro*MC9::MC9:GUS* have been described previously^35^.

To place the tonoplast integrity (ToIM) marker^6^ downstream of the *BFN1* promoter, the promoter of the putative protease *PASPA3* previously cloned upstream of the tonoplast integrity marker^6^ was replaced by the *BFN1* promoter. To do so, the *BFN1* promoter was amplified from a plasmid using primers comprising the attB4 and attB1r sequences, and recombined into pDONR P4P1r. The *PASPA3* promoter from pK7m34GW-*pPASPA3*-*eGFP*-*2A*-*sp*-*mRFP*^6^ was removed by a standard BP recombination with pDONRP4P1r, the destination vector was recovered by selecting on Chloramphenicol and Spectinomycin in DB3.1 cells and subsequently recombined during an LR reaction with pENTRL4R1-pBFN1. The resulting expression vector pK7m34GW-pBFN1-eGFP-2A-sp-mRFP was used to electroporate *Agrobacterium tumefaciens*. Agrobacterium-mediated floral dip transformation of wild-type (Col-0) plants was performed and several apparent transformants were screened for visible ToIM signal, leading to the selection of a representative pro*BFN1::ToIM* marker line.

To express the cell death agonist mammalian protein mBAX under the transcriptional control of the *BFN1* promoter, the vectors pENTRL4R1 containing the *proBFN1* fragment and pENTR221 containing the *mBAX* gene (kind gift from Moritz Nowack; unpublished) were together recombined into pK7m24GW^51^ using standard protocols for LR recombination. This vector was transferred into *Agrobacterium tumefaciens* which were used to transform by floral dip Col-0 and *ore1-2* plants. Transformants were then screened to identify at least one homozygous line in each genotype displaying *mBAX* expression.

### Confocal Laser Scanning Microscope (cLSM) analyses

3D-projections of LRP from Figure S1 were recorded as previously described^22, 52^ with a Leica SP5 cLSM in intervals of 1 h, using 488nm and 514nm laser lines sequentially to excite GFP and YFP, respectively.

All other cLSM images were acquired using a Zeiss LSM780 inverted microscope, by placing the seedlings in microscope chambers as previously described^53^. GFP and CITRINE signals (Figure 2; Figure S2) were simultaneously recorded with a spectral detector after excitation with a 488nm laser line, and separated by online fingerprinting (i.e. unmixing which occurs during, rather than after, acquisition). eGFP and mRFP signals (Movies S1, S2) from the ToIM marker^6^ were detected simultaneously upon excitation with 488nm (for eGFP) and 561nm (for mRFP) laser lines.

### Transmission electron microscopy

An early-stage LRP marker proLBD16::GFP line (a kind gift from Laszlo Bako) was used to identify regions of the root containing primordia. Tissues were fixed under vacuum in 2.5% w/v glutaraldehyde in 0.1M Na cacodylate buffer, pH7.3 for 30 min, and then without vacuum overnight at 4°C. Tissues were then stained with 1% w/v osmium tetroxide for 2 h at room temperature in the dark, washed twice in water, and then dehydrated in 15-minute steps through an ethanol series (50, 70, 90, and 100%). Root portions were then rinsed in propylene oxide and exchanged with Spurr’s resin (Polysciences) and then baked at 65°C for 2 days. Sections were mounted on formvar-coated copper grids. Contrasting was done for 45 min in 5% uranyl acetate followed by 5 min in Sato’s lead acetate staining.

### Light microscopy

All observations were performed with a Zeiss Axioplan II microscope equipped for epifluorescence microscopy.

Histochemical GUS assays were performed as described previously^35^. Seedlings were cleared and observed using differential interference contrast^18^.

LRP of seedlings harbouring the pro*BFN1::*nuc*GFP* constructs were observed while alive following mounting in liquid ½ MS medium. Both differential interference contrast and fluorescence micrographs were acquired to relate the possible presence of nucGFP signal to the LRP stage.

For TUNEL staining, seedlings were fixed for 2 h in freshly prepared 4% paraformaldehyde in phosphate buffered saline (PBS) (137 mM NaCl, 2.7 mM KCl, 8 mM Na_2_HPO_4_, 2 mM KH_2_PO_4_), and washed twice with PBS. Samples were permeabilized by 8 min incubation at room temperature in a solution containing 0.1% Triton X-100 and 0.1% trisodium citrate, followed twice by PBS wash. TUNEL labeling was performed according to manufacturer’s protocol (*In Situ* Cell Death Detection Kit, TMR red, Roche), and washed twice with PBS. Nuclei were stained by incubating samples for 10 min in the dark in 100 μl of 4′,6-diamidino-2-phenylindole (DAPI) solution (1 μg/ml in PBS), followed by three washing steps with PBS, each for 20 min.

### Lightsheet fluorescence microscopy

Seeds were sterilized for 10min with 70% ethanol supplemented with 0.05% Triton X100, washed 3 times with 70% ethanol, and incubated for 10 min with 100% ethanol before being dried out under the sterile bench. Glass capillaries (1.8 mm of diameter, Blaubrand intraMark, glass micropipettes 100 μL Ref. 7087 44) were sterilized with 70% ethanol for 20 min, then 100% ethanol for 20 min and left to dry. The capillaries were filled with ½ MS medium buffered with 0.5 g/L MES (adjusted to pH 5,8 with KOH), containing 1% (m/V) of Phytagel (Sigma Phytagel^TM^ P8169-250G) and placed in a small round plate filled with the same medium. One seed was placed on top of each capillary and stratified for 48h before transfer to long day conditions for 4 to 5 days at 22°C.

Imaging was performed on a Luxendo MuViSPIM, equipped with two-sided illumination and two cameras for detection. In the microscope, the plant was continuously illuminated (red and blue LEDs, turned off during stack acquisition) and temperature was maintained at 24°C. About 5 mm of the root surrounded by a cylinder of Phytagel medium was extruded out of the glass capillary which itself was positioned vertically in the microscope´s chamber containing liquid, sterile ½ MS medium. The shoots of the seedling were left out of the liquid medium.

Samples were excited by a 488nm laser light sheet (2.5 μm thickness at the waist) generated with Nikon Plan Fluor 10X/0.30W objectives. Laser power was kept <6 mW. The emitted fluorescence was detected by Nikon Apo 40X/0.80W DIC N2 objectives associated with a 497-553 nm band pass filter and captured using 2 Hamamatsu Orca-flash 4.0 cameras with an exposure time <100 ms. Under these conditions, both reporters (cell death marker pro*BFN1::*nuc*GFP* and plasma membrane marker pro*CASP1::CITRINE:SYP122*) were collected in the same channel. Stacks were acquired every 10 min for 24 h with a z-step of 0.250 μm spanning a total volume of 200 μm containing the growing LRP and the primary root overlying tissues.

After fusion of the two opposite views with the proprietary script of Luxendo, sample drift was corrected using the BigDataTracker plugin in ImageJ (Fiji).

## File name: Supplementary Movie 1

Description: Light sheet microscopy time-lapse imaging of the endodermal plasma membrane marker pro*CASP1*::*CIT:SYP122* and the cell death marker pro*BFN1*::nuc*GFP* around a developing LRP.

## File name: Supplementary Movie 2

Description: Light sheet microscopy time-lapse imaging of the endodermal plasma membrane marker pro*CASP1*::*CIT:SYP122* and the cell death marker pro*BFN1*::nuc*GFP* around a developing LRP (Fig. 2i). The central, elongated nuclear signal comes from the vasculature while the two nuclear signals from the right come from two overlying endodermal cells.

## File name: Supplementary Movie 3

Description: Depth-wise movie of light sheet microscopy imaging of the endodermal plasma membrane marker pro*CASP1*::*CIT:SYP122* and the cell death marker pro*BFN1*::nuc*GFP* around a developing LRP (Fig. 2i) at the time of disintegration of a marked nucleus. The presence of a LRP can be deduced from the bent shape of the overlying endodermal cells.

## File name: Supplementary Movie 4

Description: Time-lapse confocal microscopy 3D projection of the *BFN1* promoter driven tonoplast marker (pro*BFN1::ToIM*) line in *Arabidopsis* roots. The signal appearing shortly on the left-hand side comes from the vasculature while the signal from the right comes from an overlying endodermal cell.

## File name: Supplementary Movie 5

Description: Time-lapse confocal microscopy 3D projection of the *BFN1* promoter driven tonoplast marker (pro*BFN1::ToIM*) line in *Arabidopsis* roots. The signal appearing shortly on the left-hand side comes from the vasculature while the signal to the right comes from an overlying endodermal cell.

**Table S1:** List of 50 genes most correlated with BFN1 expression in the LR transcriptome. Sorted after decreasing Pearson correlation with BFN1 expression. Selected “cell death indicator genes” are highlighted in grey.

|  |  |
| --- | --- |
| AT1G11190 | BFN1 (BIFUNCTIONAL NUCLEASE I); single-stranded DNA specific endodeoxyribonuclease |
| AT5G04200 | MC9 (metacaspase 9); cysteine-type peptidase |
| AT2G44770 | ELMO/CED-12 family protein |
| AT4G18425 | DMP4 (paralog of DMP2) |
| AT1G26820 | RNS3 (RIBONUCLEASE 3); RNA binding / endoribonuclease/ ribonuclease T2 |
| AT4G17220 | ATMAP70-5 (microtubule-associated proteins 70-5); microtubule binding |
| AT4G34320 | unknown protein |
| AT1G55180 | PLDEPSILON (PHOSPHOLIPASE D ALPHA 4); phospholipase D |
| AT4G24430 | Rhamnogalacturonate lyase family protein |
| AT3G20850 | proline-rich family protein |
| AT2G14095 | EXI1 (EXITUS1) |
| AT2G37670 | WD-40 repeat family protein |
| AT4G39540 | ATSK2 (shikimate kinase 2) |
| AT1G11160 | Transducin/WD40 repeat-like superfamily protein |
| AT5G16600 | MYB43 (myb domain protein 43); DNA binding / transcription factor |
| AT5G54850 | unknown protein |
| AT3G21700 | Ras-related small GTP-binding family protein |
| AT1G63600 | Receptor-like protein kinase-related family protein |
| AT2G15320 | Leucine-rich repeat (LRR) family protein |
| AT1G01780 | LIM domain-containing protein |
| AT1G69580 | Homeodomain-like superfamily protein |
| AT3G03000 | calmodulin, putative |
| AT3G08930 | LMBR1-like membrane protein |
| AT1G09540 | MYB61 (MYB DOMAIN PROTEIN 61); DNA binding / transcription factor |
| AT2G47830 | cation efflux family protein / metal tolerance protein, putative (MTPc1) |
| AT3G23870 | unknown protein |
| AT4G28290 | unknown protein |
| AT5G58940 | CRCK1 (CALMODULIN-BINDING RECEPTOR-LIKE CYTOPLASMIC KINASE 1); ATP binding / serine/threonine kinase |
| AT1G34540 | CYP94D1; electron carrier/ heme binding / iron ion binding / monooxygenase/ oxygen binding |
| AT3G51770 | ETO1 (ETHYLENE OVERPRODUCER 1); protein binding, bridging |
| AT3G25805 | unknown protein |
| AT2G37810 | CHP-rich zinc finger protein, putative |
| AT5G13990 | ATEXO70C2 (exocyst subunit EXO70 family protein C2); protein binding |
| AT1G60890 | phosphatidylinositol-4-phosphate 5-kinase family protein |
| AT1G61950 | CPK19; ATP binding / calcium ion binding / calmodulin-dependent protein kinase / serine/threonine kinase |
| AT5G15600 | SP1L4 (SPIRAL1-LIKE4) |
| AT1G05310 | pectinesterase family protein |
| AT4G10370 | DC1 domain-containing protein |
| AT1G54060 | ASIL1 (ARABIDOPSIS 6B-INTERACTING PROTEIN 1-LIKE 1); sequence-specific DNA binding / transcription factor |
| AT3G14640 | CYP72A10; electron carrier/ heme binding / iron ion binding / monooxygenase/ oxygen binding |
| AT4G01110 | unknown protein |
| AT4G09890 | unknown protein |
| AT1G01140 | CIPK9 (CBL-INTERACTING PROTEIN KINASE 9); ATP binding / kinase/ protein kinase/ protein serine/threonine kinase |
| AT4G17170 | RABB1C (ARABIDOPSIS RAB GTPASE HOMOLOG B1C); GTP binding / GTPase |
| AT5G03640 | protein kinase family protein |
| AT2G11150 | transposable element gene |
| AT4G18430 | AtRABA1e (Arabidopsis Rab GTPase homolog A1e); GTP binding |
| AT2G20880 | AP2 domain-containing transcription factor, putative |
| AT5G44700 | GSO2 (GASSHO 2); ATP binding / protein binding / protein kinase/ protein serine/threonine kinase/ protein tyrosine kinase |
| AT5G47250 | disease resistance protein (CC-NBS-LRR class), putative |
| AT4G35190 | LOG5 (LONELY GUY 5); putative lysine decarboxylase family protein |

**Figure S1:**
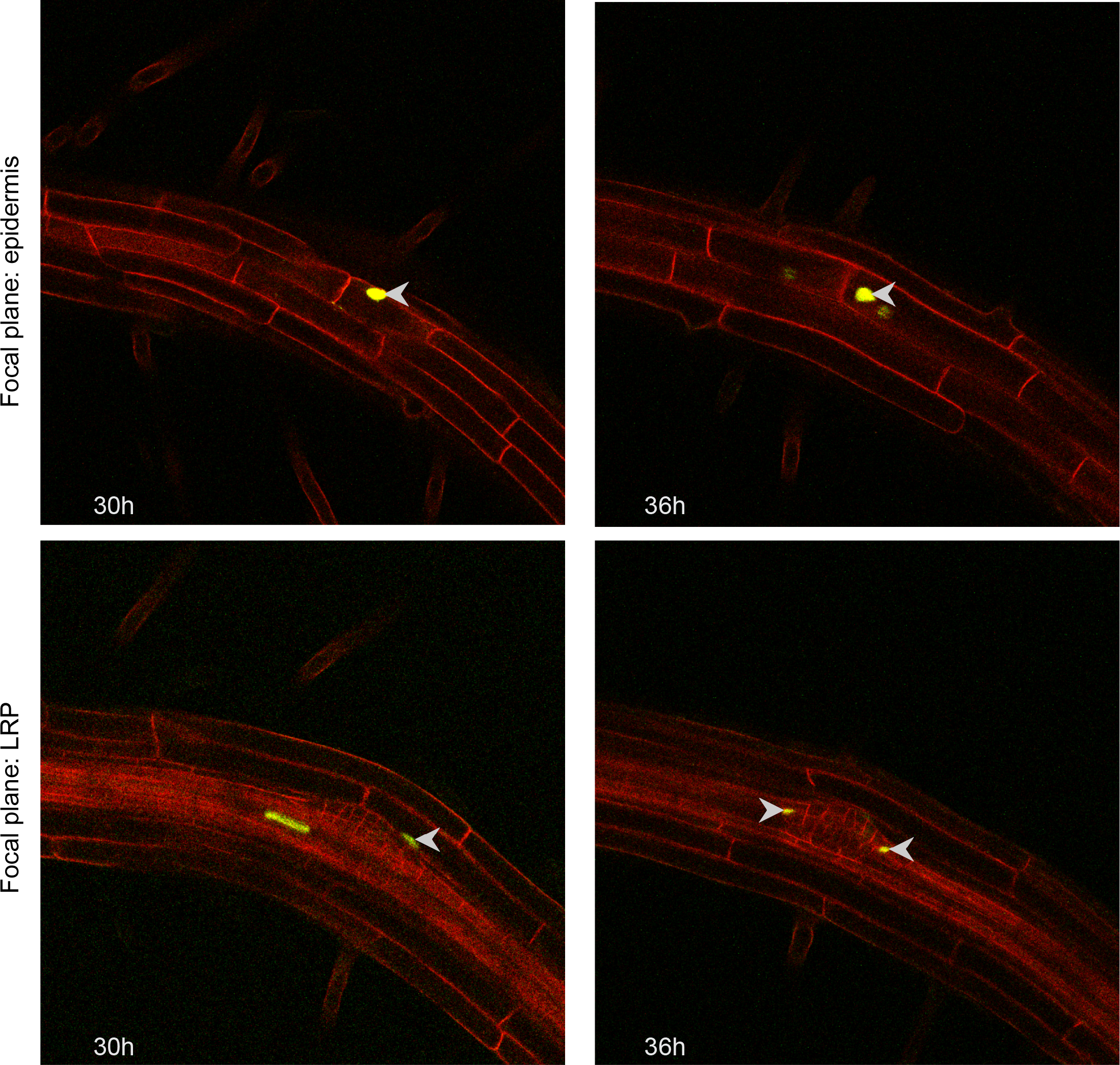
Vizualization of proBFN1 activation in cells overlying early-stage LRP. pro*BFN1::*nuc*GFP* expression at two time points during lateral root emergence (LRE) of a gravitationally induced lateral root (LR) in pro*UBQ10::WAVE131:YFP* (ubiquitous plasma-membrane marker) background. Arrowheads indicate GFP-positive nuclei in overlying cells, while the elongated fluorescent nucleus in the stele belongs to a xylem tracheary element, a cell type known to undergo cell death.

**Figure S2:**
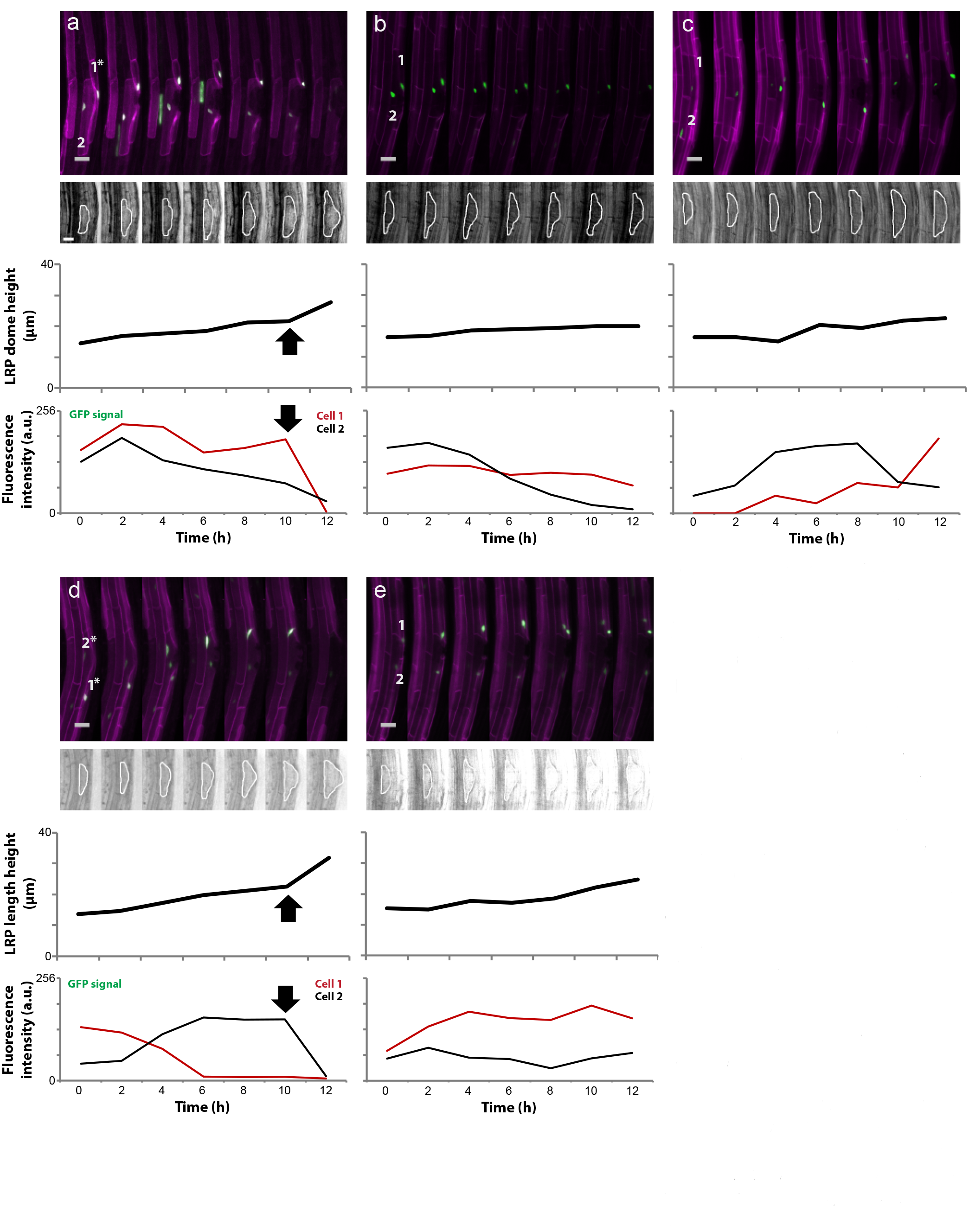
Viability of LRP-overlying cells in relation to LRP length. (a-e) Time-lapse confocal microscopy observation of five different 6-days-old Arabidopsis seedlings harboring an endodermal-specific plasma-membrane marker (*proCASP1:: CITRINE:SYP122*) and a nuclear marker driven by a cell death indicator gene’s promoter (*proBFN1::nGFP*). Upper micrographs represent 3D projections of fluorescence signal around a growing LRP. Lower micrographs display transmission light images of the corresponding LRP, underlined by a white line. Upper charts show the height of each LRP. (a) is reproduced from Fig. 2h. Lower charts display quantifications of the nuclear GFP fluorescence intensity in the cells marked by numbers in the upper micrographs. Asterisks indicate endodermal cells which die during LRP progression. Scale bars = 25μm. Black arrows mark accelerations in LRP growth concommitant with overlying endodermal cell death.

**Figure S3:**
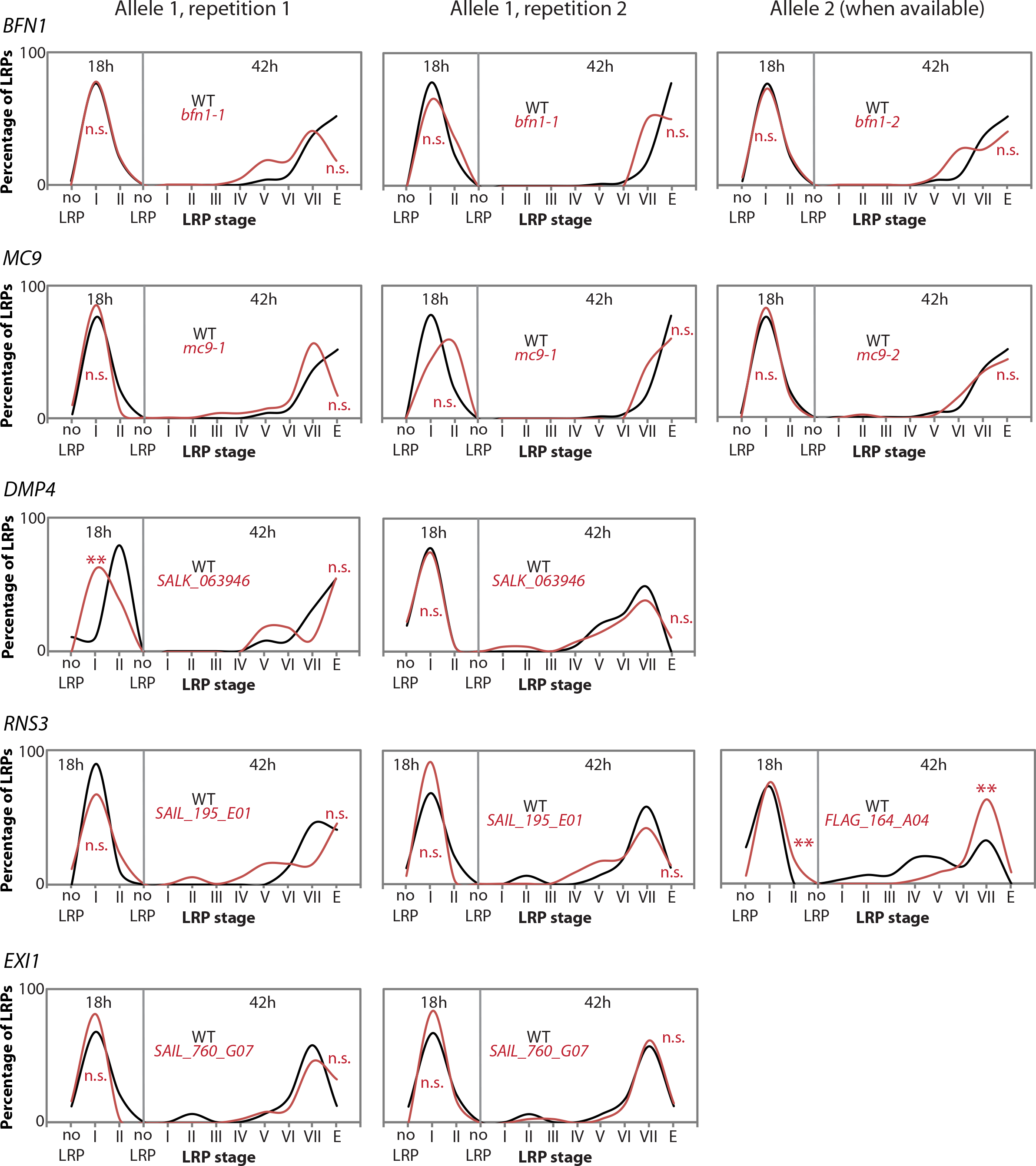
Staging assay on T-DNA insertion lines for cell death indicator genes. Gravitrational induction of LRP initiation was applied to T-DNA insertion lines for the cell death indicator genes. The stage of LRP development was monitored for seedlings of each genotype (including control wild-type) 18h and 42h after gravitropic induction, allowing to display the proportion of LRPs at each stage at these time-points. Each T-DNA insertion line was compared to the corresponding wild-type using Pearson’s chi-square test to reveal potential differences in the distribution of LRP stages dependent on genotype (n.s.: not significant, **: p<0.01).

**Figure S4:**
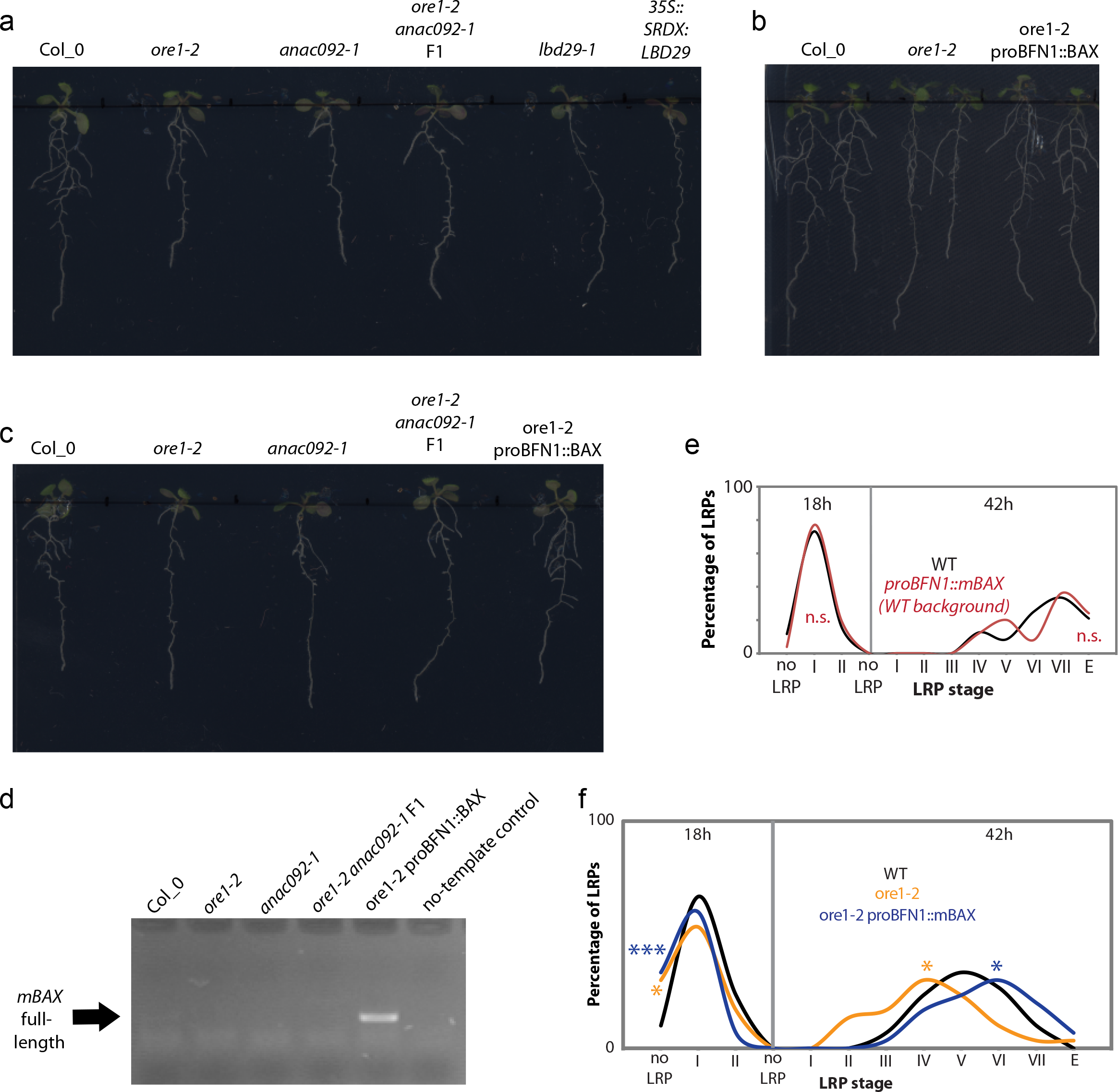
Characterization and rescue of the lateral root emergence phenotype in mutants for the positive cell death regulator ORE1. (a-c) Images of seedlings illustrate the differences in the numbers of emerged lateral roots (LRs) between the indicated genotypes grown next to one-another (i.e. in the same conditions). (d) The gel image displays an agarose gel electrophoresis with products from PCRs with cDNA from the indicated genotypes and designed to amplify the mamalian *mBAX* transcript, which has been introduced in the *ore1-2* mutant background. (e) The chart shows the results of a staging assay comparing LR emergence between wild-type plants (WT) and plants expressing the cell death inducing mammalian protein mBAX under the transcriptional control of the *BFN1* promoter (in the WT background). n.s. indicates that there is no statistically significant difference. (f) Distribution of observed LRP stages 18h and 42h after gravitational induction of LR initiation in wild-type, *ore1-2* mutant and *ore1-2* expressing the cell death inducing mBAX protein under the transcriptional control of the *BFN1* promoter. Each line was compared with the corresponding wild-type by Pearson’s chi-square test, to reveal potential differences in the distribution of LRP stages dependent on genotype (n.s.: not significant, *: p<0.05, **: p<0.01, ***:p<0,001). n=30 observed seedlings per genotype and per time-point.

**Figure S5:**
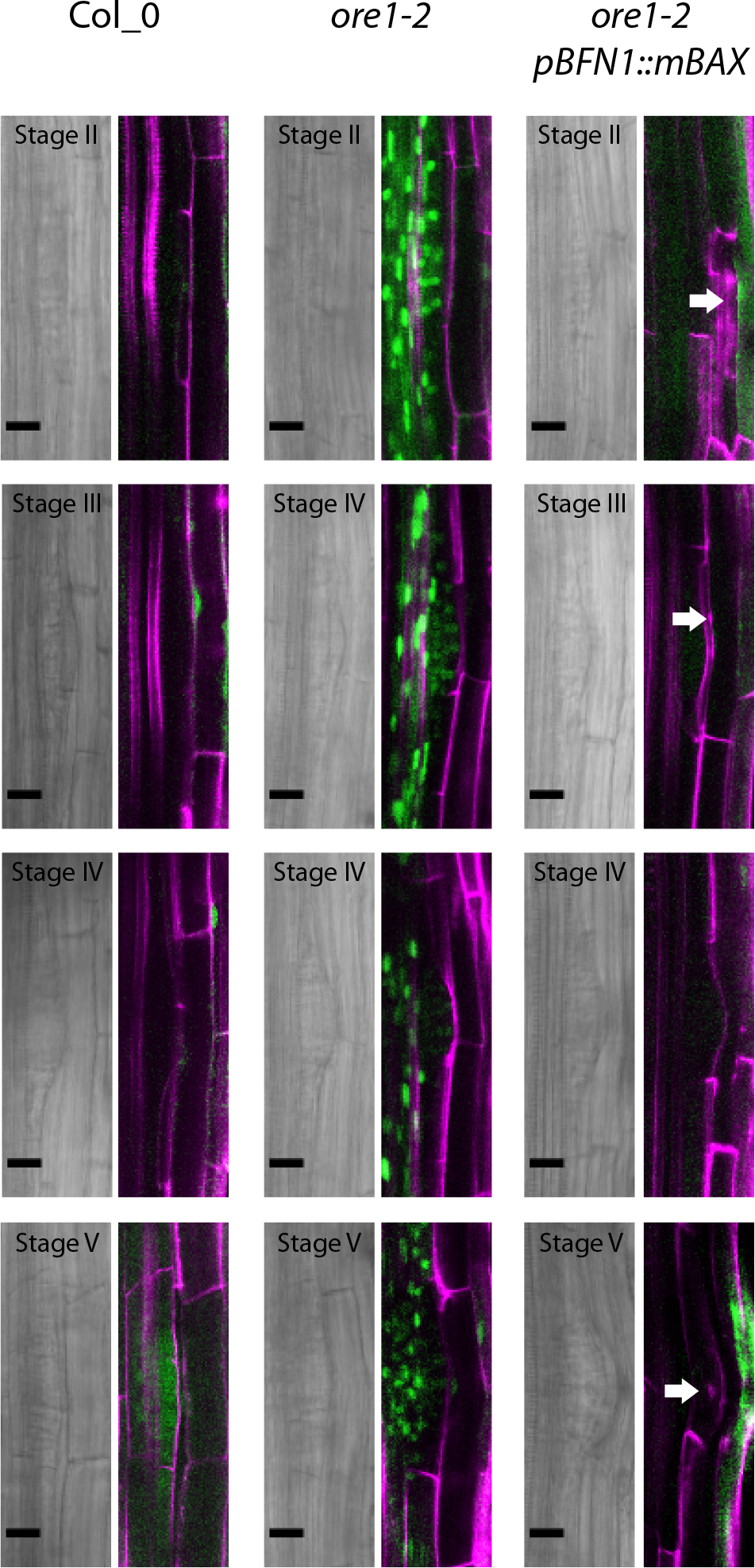
Evidence for mBAX-induced cell death in LRP overlying cells. Transmission light (left) and fuorescence (right) micrographs from confocal laser scanning microscopy imaging of LRPs and of the overlying cells in the main roots of 6 days old *Arabidopsis* seedlings. The seedlings were stained with propidium iodide (magenta) and fluorescein diacetate (FDA; viability staining; green). White arrows point at cells displaying signs of cell death and autolysis, as revealed by complete lack of FDA signal as well as by propidium iodide staining of the nucleus or of collapsed protoplast remains.

